# A heritable androgenic mechanism of female intrasexual competition in cooperatively breeding meerkats

**DOI:** 10.1101/2021.01.11.425748

**Authors:** Christine M. Drea, Charli S. Davies, Lydia K. Greene, Jessica Mitchell, Dimitri V. Blondel, Caroline L. Shearer, Joseph T. Feldblum, Kristin A. Dimac-Stohl, Kendra N. Smyth-Kabay, Tim H. Clutton-Brock

## Abstract

Female intrasexual competition can be intense in cooperatively breeding species, with the dominant breeder or matriarch limiting reproduction in subordinates via aggression, eviction or infanticide. In males, these tendencies bidirectionally link to testosterone, but in females, there has been no systematic investigation of androgen-mediated behaviour within and across generations. In 22 wild meerkat (*Suricata suricatta*) clans, we show that matriarchs 1) express peak androgen concentrations during late gestation, 2) when displaying peak feeding competition, dominance, and evictions, and 3) relative to subordinates, produce offspring that are more aggressive in early development. Late-gestation, antiandrogen treatment of matriarchs 4) reduced their dominance behaviour, was associated with infrequent evictions, decreased social centrality within the clan, 5) increased aggression in cohabiting subordinate dams, and 6) reduced their offspring’s aggression. These effects implicate androgen-mediated aggression in the operation of female sexual selection, and intergenerational transmission of ‘masculinised’ phenotypes in the evolution of meerkat cooperative breeding.

## Introduction

In cooperatively breeding species, nonbreeding helpers assist in raising young born to dominant breeders, such that both sexes experience pronounced reproductive skew. Whereas the functional explanations for why helpers forfeit reproduction to assist others implicate benefits gained through kin selection or inclusive fitness (augmented through mutualism or reciprocity) (*1, 2*), the proximate physiological and behavioural mechanisms of reproductive monopoly or suppression can be elusive (*3, 4*), particularly if supporting evidence is strictly correlational. Studies on the neuroendocrine mediators of cooperative breeding are often focused on *reproductive suppression*, targeting the role of oxytocin, vasopressin, prolactin and testosterone in alloparental care (*5, 6*) or the role of glucocorticoids, progestins and oestrogens in subordinate infertility (*7, 8*). Less is known about the neuroendocrine mediators of *reproductive monopoly*.

Whereas androgen-mediated power struggles over reproductive opportunities, including infanticide, are commonplace among male mammals (*9–11*), the same is uncharacteristic of their female counterparts (*12*). Nevertheless, in some species (*13, 14*), including cooperative breeders (*15–19*), females express raised androgen concentrations relative to conspecific males, that relate to female dominance or breeding status. Although typically adversative to females (*20*), androgens may be key to explaining female reproductive advantage under certain circumstances, specifically via aggressively mediated, female intrasexual competition. We examine this possibility in the meerkat (*Suricata suricatta*).

The meerkat is a social mongoose and obligate cooperative breeder that displays a despotic, rather than linear, hierarchy: A single dominant pair, particularly the matriarch, largely monopolises aseasonal, asynchronous reproduction within a clan of up to 50 subordinate individuals (*15*). Unlike many other cooperative breeders, meerkats adhere to a ‘limited control’ model of reproductive skew, as all adults are physiologically capable of breeding. Even if the degree of reproductive skew varies socially and environmentally (*21*; Supplementary Material, Fig. S1), reproductive success depends on competitiveness (*15, 22*). Any dam can commit infanticide, but evictions of subordinate females are typically performed by the dominant matriarch. Status-related differences in concentrations of glucocorticoids, progestins and oestrogens exist (*21–24*), but cannot explain female reproductive ‘suppression’ (*21*). Instead, the highly aggressive nature of female meerkats (*15, 25*), coupled with their exceptionally high androgen concentrations, particularly whilst pregnant (*19*), raise the possibility of a novel means of female reproductive competition within a cooperatively breeding mammal: We propose that female meerkats are naturally physiologically and behaviourally ‘masculinised’ (sensu *26*) by the actions of their own androgens, with dominant and subordinate females being masculinised to different degrees, potentially differentiating their daughters’ respective reproductive trajectories.

Two modes of hormone action would contribute to this phenomenon: Concurrent and reversible *activational* effects of late-to-post gestational androgens could facilitate the matriarch’s aggression at an opportune time, both for expelling reproductive competitors (when access to resources is critical) and for protecting vulnerable offspring (when threat of infanticide is high; *27*). Concomitantly, reflecting the temporal prenatal sequence in mammalian sexual differentiation (from gonadal, to internal then external genitalic, to neurological) (*28*), permanent and delayed *organizational* effects of key late-term, maternal androgens (*29–31*) on the developing neural circuitry of foetal daughters could promote differences in their future aggressiveness, based on maternal status, without interfering with reproductive function (*29, 32*). As may be the case in select species (*14, 30, 31, 33*), daughters from different mothers might be differentially advantaged to later compete for dominant status and, ultimately, gain reproductive success. Here, we combine normative and experimental data to, respectively, ascertain female phenotypes and test for both activational and organizational behavioural effects that would reveal a heritable masculinising mechanism of female meerkat breeding competition.

The ‘female masculinisation hypothesis’ leads to three initial predictions about normative, status-related and temporal patterns in meerkat behavioural endocrinology: In addition to 1a) greater androgen concentrations in dominant than subordinate dams (*15, 19*), we expect 1b) androstenedione and testosterone concentrations to increase across meerkat gestation and potentially into the early postpartum period (*13, 34*), 2a) a corresponding rise in the matriarchs’ competitive behaviour (directed at any clan member) and 2b) in the prosocial or submissive behaviour she receives from subordinates, as well as 3) an organizational effect of differential, late-term androgen exposure on offspring aggression. The latter should be evident in offspring of both sexes, before the onset of activational effects at puberty, namely in the behaviour of pups (1-3 months) and juveniles (3-6 months) (*31*).

An experimental test of this hypothesis then requires either eliminating androgens or interfering with their actions (e.g. via androgen receptor blockade), specifically in dominant dams late in gestation and into the early postpartum period. Such a manipulation produces an additional three predictions: Treated matriarchs should 4a) concurrently diminish their competitive and dominance-related behaviour, 4b) potentially altering social dynamics within the clan, as evidenced by reduced social centrality, 5) concurrently increase competition between or diminish submission in cohabiting subordinate dams and, 6) ultimately, bear offspring that show altered behaviour, specifically reduced prepubertal aggression relative to the offspring of control matriarchs.

## Results

### Prediction 1: Female gestational androgens should differ by status and increase across pregnancy

Studying 22 well-habituated meerkat clans in the Kalahari Desert of South Africa (*15*; tables 1–3), we found our predictions about normative patterns to be supported. First, consistent with prediction 1a, androgen concentrations in dominant females, during and post pregnancy, exceeded those in subordinate females (Generalised Linear Mixed Models (GLMMs): androstenedione *χ*^2^_1,31_ = 50.49, *P*< 0.001; testosterone *χ*^2^_1,32_ = 6.39, *P* = 0.011; Linear Mixed Model (LMM): faecal androgen metabolites (fAm) *χ*^2^_1,7_ = 7.60, *P* = 0.006; Fig. 1A-C; table S1). Additionally, consistent with prediction 1b), overall female androgen concentrations changed significantly from early, mid, to late pregnancy (EP, MP and LP, respectively) and to early postpartum (PP), defined as the first two weeks after birth (GLMM: androstenedione *χ*^2^_2,12_ = 9.50, *P* = 0.008; testosterone *χ*^2^_2,11_ = 9.50, *P* = 0.009; LMM: fAm *χ*^2^_2,7_ = 7.19, *P* = 0.027), reaching their peak during LP (Least Significant Difference (LSD): androstenedione LP > MP, *t* = −2.86, *P* = 0.035; testosterone LP > EP, *t* = −2.94, *P* = 0.033; fAm LP > PP, *t* = 2.61, *P* = 0.027; Fig. 1A-C; table S1). There were no significant interactions between status and pregnancy stage, indicating that dominant females maintained their androgen ‘advantage’ over subordinates consistently across gestation.

**Fig. 1.**
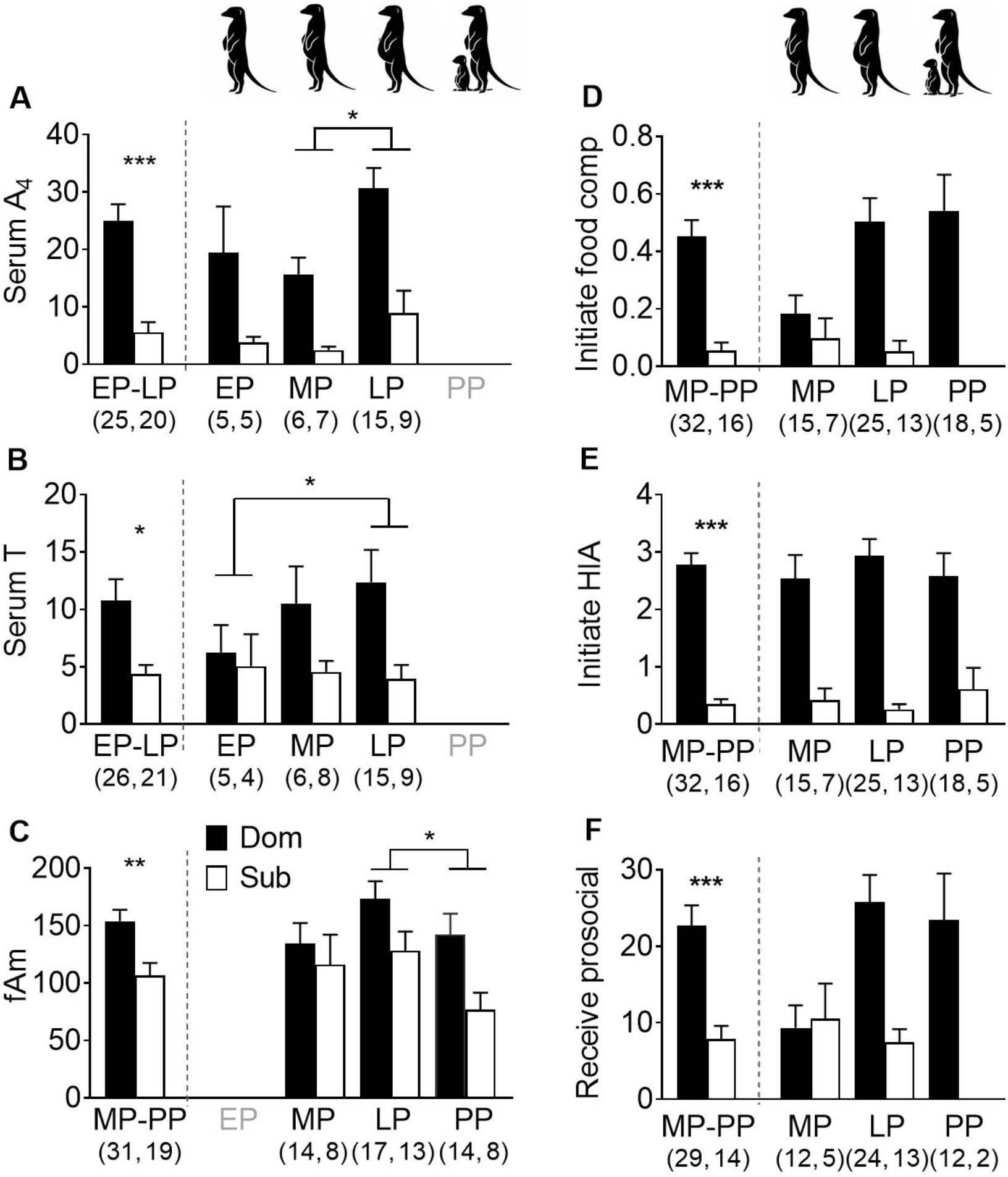
Status-related differences in androgen concentrations and behavioural rates during and after pregnancy in wild meerkats. Mean (+ S.E.) concentrations of (**A**) androstenedione (A4, ng/ml), (**B**) testosterone (T, ng/ml), (**C**) faecal androgen metabolites (fAm, ng/g), and hourly rates of (**D**) initiated food competition (a subset of high-intensity aggression, ‘HIA,’ specific to foraging and salient to reproductive competition), (**E**) all initiated HIA throughout the day, and (**F**) received prosociality at the den, both overall (pre dashed lines) and by early, mid and late pregnancy (EP, MP and LP, respectively), and the first two weeks postpartum (PP; post dashed lines). Values in parentheses represent the numbers of (**A-C**) pregnancies and (**D-F**) females sampled at different locales, by dam status: dominant (Dom, black) and subordinate (Sub, white). *** *P* < 0.001, ** *P* < 0.01, * *P* < 0.05. (Icons by S. Bornbusch.)

### Prediction 2: Female aggression during pregnancy should differ by status and increase across gestation

In the Kalahari, where resources are limited, food competition (a subset of intense aggressive behaviour) is particularly salient to meerkat reproductive competition, as are aggressively mediated evictions of reproductive competitors. Consistent with prediction 2a, dominant dams more frequently initiated food competition (GLMMs: χ^2^_1_ = 13.45, *P* < 0.001; Fig. 1D), initiated more intense overall aggression (χ^2^_1_ = 35.39, *P* < 0.001; Fig. 1E) and engaged in more ‘status-advertising’ scent marking (χ^2^_1_ = 10.81, *P* = 0.021; table S2) than did subordinate dams across reproductive stages. Likewise (prediction 2b), dominant dams also received more prosociality from clan members than did subordinate dams (χ^2^_1_ = 14.73, *P* < 0.001; Fig. 1F). When only matriarchs were included in the time-course analysis, rates of initiating food competition (χ^2^_2_ = 9.78, *P* = 0.008) and receiving prosociality (χ^2^_2_ = 18.80, *P* < 0.001; LSD: LP > MP *t* = −3.14, *P* = 0. 004) increased significantly across pregnancy. The matriarch’s androgen concentrations (Fig. 1A-C) and dominance interactions (Fig. 1D, F) thus peaked in LP, when the evictions of subordinate females, inferred from day-to-day changes in clan composition, also occur most frequently (*27*).

### Prediction 3: Maternal status should differentiate normative infant aggression

Regarding maternal effects, and consistent with prediction 3, analysis of meerkat behavioural ontogeny revealed powerful, and heretofore unknown, patterns in early offspring aggression. These patterns are suggestive of organizational effects of androgens, as the offspring (from both sets of mothers) showed no sex difference (i.e., females were just as aggressive as males; table S3) and those from dominant, relative to subordinate, mothers initiated more aggression throughout the first six months of life (Fig. 2; table S3). Moreover, pup aggression started high, coincident with peak competition for attention from and feeding by helpers, then declined with advancing age to juvenility (GLMM: χ^2^_1_ = 255.98, *P* < 0.001; table S3), alongside the acquisition of independent feeding. Reflecting the different initial rates of aggression, there was a steeper negative slope in offspring from dominant versus subordinate mothers (LSD: *t* = 2.87, *P* = 0.011; Fig. 2; table S3).

**Fig. 2.**
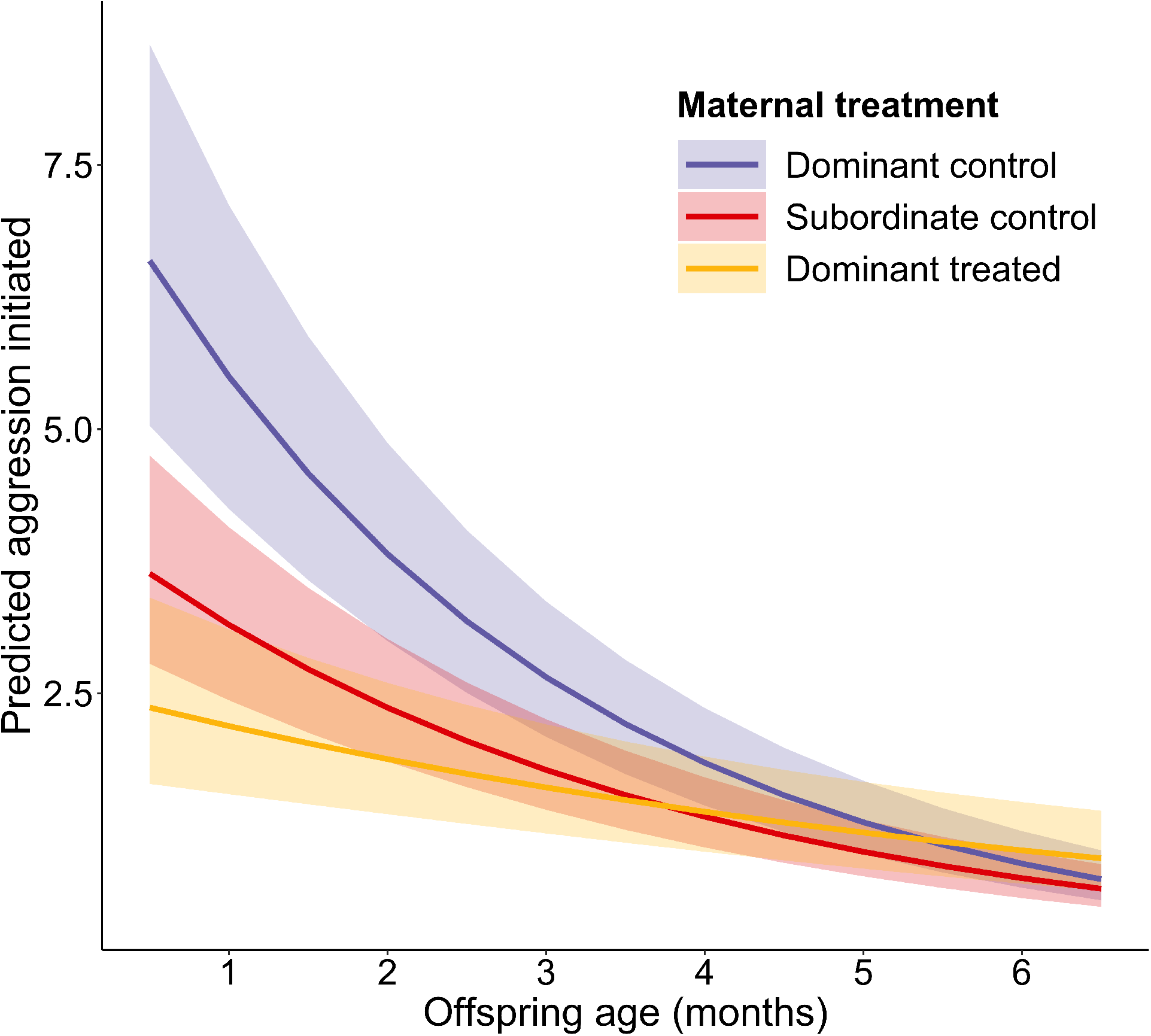
Different model-predicted rates of offspring aggression by age, based on maternal status and treatment condition. The mean predicted rates (frequency/mean focal duration or 14.4 min, with 95% confidence intervals) of initiated aggression by offspring during their first six months of life (pups: 1-3 mo; juveniles: 3-6 mo), are shown with respect to maternal status and antiandrogen treatment: dominant control (purple), subordinate control (red), and dominant treated (orange) dams.

### Prediction 4: Experimental blockade of androgen receptors in matriarchs should alter their concurrent, androgen-mediated behaviour

To experimentally test the ‘female masculinisation hypothesis,’ we used previously validated methods (*35*) to administer the antiandrogen, flutamide, to a subset of dominant dams late in gestation and into the early postpartum period (Fig. S2). In line with the variable effects of flutamide on circulating androgen concentrations across species, including male meerkats (*38*), treatment of female meerkats had no appreciable effects on their concentrations of faecal androgen metabolites across the full, three-week treatment period (Fig. S3). When accounting for flutamide-associated decreases in meerkat aggression, the potential role of aromatization of androgens to oestrogens could not be discounted in males (given lack of knowledge about male oestrogen concentrations and their behavioural correlates; *38*); however, the role of aromatization can be discounted in dominant female meerkats, given the existing information on female oestrogen concentrations (*19, 21–23*). Thus, when androgen receptors were blocked, specifically prohibiting the actions of androgens, our predictions about experimentally induced patterns in females were also supported.

Consistent with prediction 4a, during LP-PP and relative to control matriarchs, treated matriarchs showed significantly reduced rates of initiating food competition (GLMMs: χ^2^_1_ = 7.39, *P* = 0.007; Fig. 3A), scent marking (χ^2^_1_ = 6.94, *P* = 0.008; Fig. 3B), and receiving prosocial (χ^2^_1_ = 5.73, *P* = 0.017; Fig. 3C) and submissive (χ^2^_1_ = 4.26, *P* = 0.039; Fig. 3D) behaviour from clan members (table S4). Thus, whether owing directly to their reduced aggression or indirectly to their infrequent or potentially altered scent signals, treated matriarchs commanded less ‘attention’ from subordinate clan members than did control matriarchs. In addition, relative to control matriarchs, treated matriarchs evicted nearly two-thirds fewer subordinate females (Fig. 3E), but so few evictions precluded statistical modeling.

**Fig. 3.**
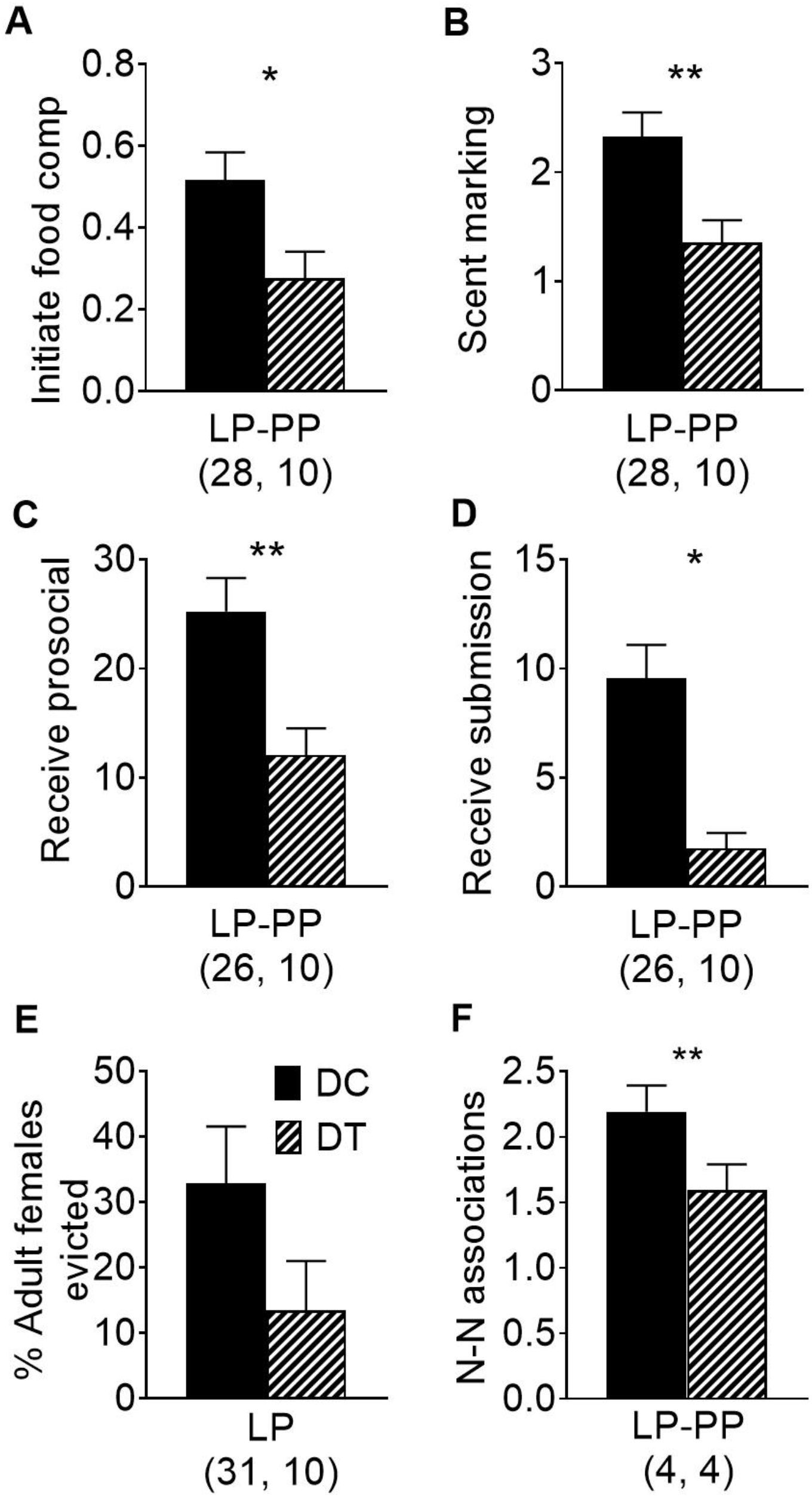
Effects of antiandrogen treatment on the behaviour of dominant dams. Shown for dominant control (DC, black) versus dominant treated (DT, bold hatching) dams are the matriarchs’ mean (+ S.E.) hourly rates of (**A**) initiated food competition (as in Fig. 1), (**B**) scent marking, (**C**) received prosociality, and (**D**) received submission during late pregnancy (LP) and early postpartum (PP) for dominant control (DC, black) and dominant treated (DT, bold hatching) dams. Also shown are the mean (+ S.E.) (**E**) percentages of adult females evicted during LP and (**F**) hourly rates (scans/partner/hr) of dyadic nearest-neighbour (N-N) associations with adult clan members during LP and PP. In (**F**), matriarchs were matched across control and treated conditions in four clans. Whilst treated, these four dams were in close proximity to another adult clan member during ~1 fewer scan/individual/2 hrs observation. For a clan with a mean of 14 adults, this value corresponds to a predicted difference of ~8 N-N associations/hr. Values in parentheses represent the numbers of (**A-E**) pregnancies or (**F**) matched clans sampled. ** *P* < 0.01, * *P* < 0.05.

Consistent with prediction 4b, in a subset of four clans in which each of the four matriarchs experienced, in random order, both a control and a treated pregnancy (such that respective clan sizes, territories, memberships and social dynamics were matched across randomised conditions), antiandrogens significantly reduced the frequency of associations between the matriarch and adult clan members (LMM, *t*51.15 = −3.04, *P* = 0.004; Fig. 3F; table S5). Thus, within a mere three-week treatment window, treated dams behaved in a manner less characteristic of matriarchs, received less deference from clan members, appeared to evict fewer females and became less socially central within their clan.

### Prediction 5: Effects of experimental manipulation in matriarchs should influence the behaviour of subordinate dams

Concurrently pregnant subordinates represent a threat to the matriarch’s reproductive interests, as their pups can dilute the benefits provided by helpers. Because subordinate dams face an increased risk of being evicted, they typically maintain a low profile within the clan. It is notable, however, that dominance overthrows are often preceded by increased competition between subordinate contenders. Consistent with prediction 5, when the pregnant matriarch received antiandrogens, concurrently pregnant subordinates within her clan initiated significantly more food competition (GLMM: χ^2^_1_ = 6.48, *P* = 0.011; Fig. 4A) and intense aggression (χ^2^_1_ = 6.55, *P* = 0.010: Fig. 4B) whilst foraging than did subordinate dams whose pregnant matriarch was left untreated (table S6). The subordinate dams’ initiation of prosocial and submissive behaviour at the den also appeared to decrease (Figs. 4C, D), but the short periods spent above ground, at the den (versus whilst foraging), produced smaller sample sizes that prevented model convergence. Overall, pregnant subordinates, specifically, responded to the matriarch’s treatment in a manner that could potentially improve their access to resources, the defence of their own offspring and their threat to the matriarch’s offspring.

**Figure 4.**
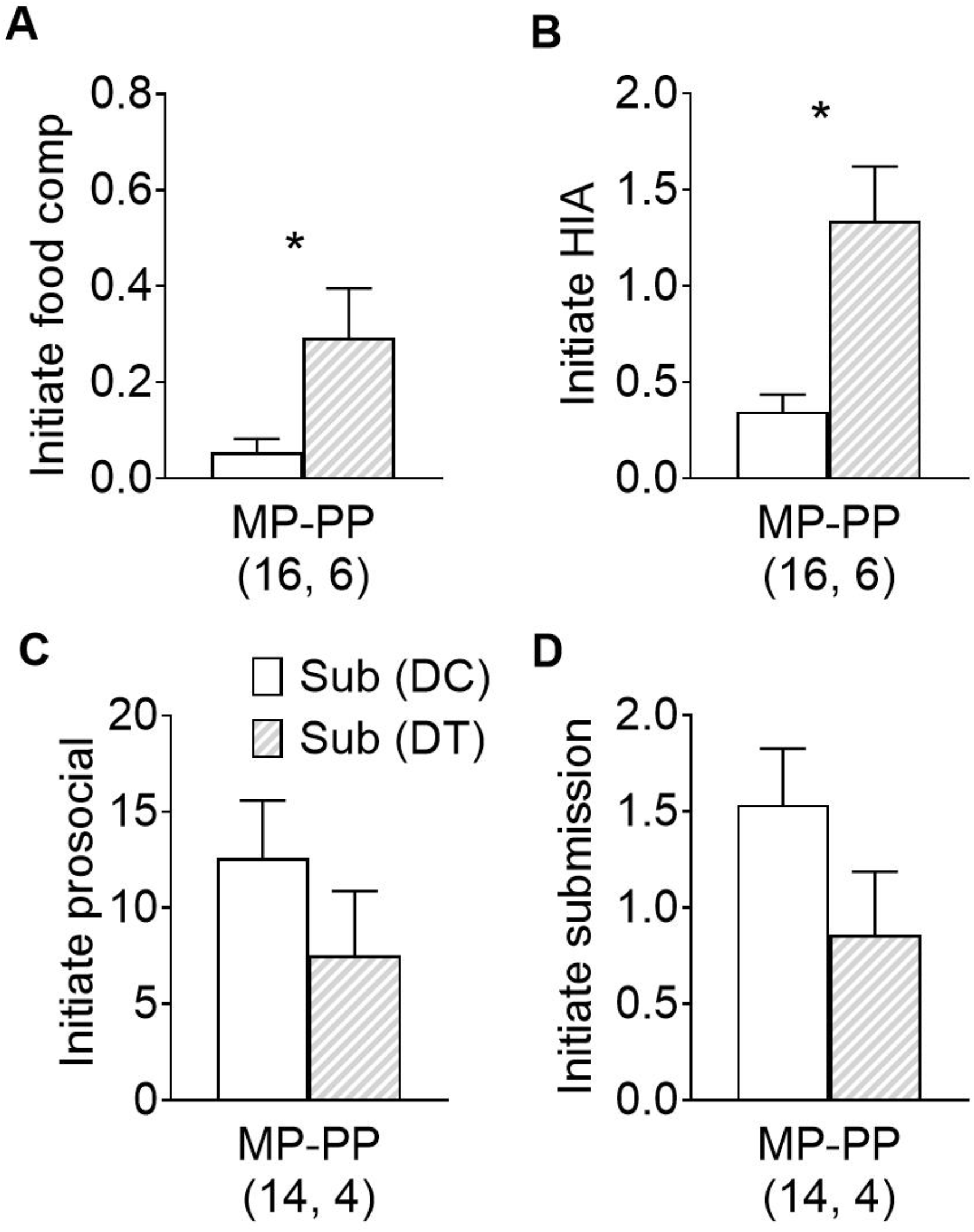
Effects of antiandrogen treatment of dominant dams on the behaviour of concurrently pregnant subordinates within the same clan. Shown are mean (+ S.E.) hourly rates of initiated (**A**) food competition, (**B**) high-intensity aggression, HIA, (**C**) prosociality and (**D**) submission during mid pregnancy (MP), LP and/or PP for cohabiting subordinate dams under their matriarch’s regimens (Sub DC, white; Sub DT, light hatching). The subordinates’ broader range of pregnancy stages (relative to Fig. 4) accommodates variability in the synchrony of concurrent pregnancies. Values in parentheses represent the numbers of pregnancies sampled. * *P* < 0.05.

### Prediction 6: Blocking the actions of maternal androgens in late gestation should decrease infant aggression

Lastly, evidence of ‘carry-over’ effects of maternal treatment on pups represent the ultimate test of organizational effects. Consistent with prediction 6, the offspring of antiandrogen-treated matriarchs, regardless of sex (table S3), initiated significantly less aggression than did those of control matriarchs throughout the first six months of life (Fig. 2; table S3). The age-related decline in aggression by offspring from control matriarchs was greatest relative to that by offspring from treated matriarchs (LSD: *t* = 2.34, *P* = 0.049; Fig. 2; table S3). These findings confirm that maternal androgens normally reach foetal meerkats in LP and PP, presumably via the placenta and maternal milk, respectively, and influence later expression of their behavioural phenotype. Indeed, whilst experimental ‘elimination’ of the natural, status-related difference in intergenerational patterns (i.e., similar aggression rates between offspring from treated matriarchs and subordinate controls; LSD: *t* = −0.12, *P* = 0.992; Fig. 2) also confirms the role of androgens in normative behavioural patterns, it further indicates physiological relevance of the antiandrogen dosage and treatment duration.

## Discussion

This integrated, intergenerational study presents multiple elements, both from normative and experimental data sets, to support our unified hypothesis about female masculinisation in the cooperatively breeding meerkat. Specifically, androgenic action in females explains both the natural, status-related differences in adult aggression and a maternal or transgenerational effect on offspring aggression. The supporting elements from the mother’s generation include the natural physiological and behavioural differences between dominant and subordinate dams, the direct mirroring in flutamide-treated matriarchs of the natural, ‘subordinate’ behavioural phenotype and the indirect effects of their treatment on the behaviour of pregnant subordinate clan members. The immediacy of adult behavioural effects from androgen receptor blockade are consistent with those observed in treated male conspecifics (*35*) and reversed from those observed in androgen-supplemented females of other species (*36*). We suggest that over a longer time-course than three weeks, antiandrogen treatment could potentially contribute to an overthrow of female dominance relations, breeding status and even reproductive outcomes.

The supporting elements from the offspring’s generation include the natural ‘inheritance’ of maternal behavioural phenotypes and the mirroring of ‘subordinate’ behaviour by the descendants of treated matriarchs, evident prepubertally through to six months of age. These transgenerational findings are consistent with female behavioural masculinisation in all lineages, but with dams of differing status exerting different levels of hormonal influence over their respective pups (as in *14*). Moreover, these findings reflect consistency in the reduced expression of androgen-mediated traits via flutamide treatment, as seen in male meerkats and in the highly masculinised, female spotted hyaena (*Crocuta crocuta*) (*37, 38*).

Unlike the typical, mammalian developmental pattern of early prosociality, followed by increasing, male-biased aggression, our findings on the absence of offspring sex differences and the early emergence of aggression are also consistent with patterns observed in other behaviourally masculinised mammals (*14, 31, 39*). Indeed, spotted hyaenas show the same unusual temporal pattern (*39*) and, based on correlational data, the same intergenerational pattern (*14*). Nevertheless, because maternal, behavioural interventions ensure status acquisition of offspring in hyaena society (*40*), hyaena mothers help shape their cubs’ aggressive interactions; by contrast, the cooperative rearing of meerkat pups minimizes the influence of certain maternal factors (*41*), limiting the potential for a similar socialisation confound. The meerkat system thus better isolates the influence of biologically relevant differences in prenatal androgens on early aggression. These data illustrate the different potentials for androgen-mediated, female aggression in different meerkat ‘lineages’ – potentials that are activated in adulthood and organized *in utero*.

The late-gestation timing of peak androgen exposure is relevant both to activational effects on mothers, that enable protection of their reproductive investment, and to organizational effects on their developing daughters, when foetal development is most sensitive to sexual differentiation of the embryonic neural structures underlying postnatal behaviour. Timing in relation to the endocrine milieu matters because despite clear physiological masculinisation (*19*; Fig. 1A-C), the reproductive organs of female meerkats are not prenatally masculinised, as has been observed in other exceptional species (*42–45*). Although various androgenic mechanisms can account for genital masculinisation of foetal females (*43, 46*), differential genital versus behavioural masculinisation (e.g. *44*) traditionally owes to the timing, quantity and duration of foetal androgen exposure (*32*) and, potentially, to the type of androgen (i.e., androstenedione versus testosterone; *29, 31*). With regard to timing, developing daughters can be protected from reproductive consequences of early-to mid-gestational androgens, for example, via selective enzymatic action in utero (e.g. aromatase enhancing early placental conversion of maternal androgens to oestrogens: *47*). Evidently protected from potential costs of early masculinisation (e.g. *20*), meerkats, like other vertebrate females (*48, 49*), nonetheless experience costs of natural androgens (again, potentially differentiating androstenedione from testosterone), including increased parasitism (*50*) and reduced immunocompetence (*51*). Although androgen-mediated health costs may carry reproductive tradeoffs during environmental stressors (*21*), the socio-reproductive advantages appear to far outweigh these costs.

Across mammals, we specifically lack understanding of the mechanisms underlying female social dominance and pronounced aggression, both at an individual and species level. The documented importance of androgens in the natural behavioural repertoire of female meerkats addresses this gap and suggests a broader role for androgens in female mammals than previously recognized. Whereas activational effects of androgens are temporary, organizational effects typically endure for the lifetime of the affected individual, even if expression may become latent (such as when meerkat offspring settle into uniformly submissive roles as juveniles). Thus, beyond accounting for an uncharacteristically aggressive female phenotype, androgen concentrations differentiate dominant and subordinate females and differentially affect the aggressive potential of their offspring – potential that daughters can call upon in adulthood to improve dominance acquisition and maintenance, and, ultimately, reproductive success. Androgens thus appear to underly female sexual selection in meerkats.

As a final note, although maternal effects can be ‘inherited’ by daughters, they may not necessarily transmit to subsequent generations. A mechanism to perpetuate physiological masculinisation may owe to an additional androgen-mediated, transgenerational, epigenetic mechanism operating during female sexual differentiation (*52*). Such a mechanism has been identified in the highly masculinised, female spotted hyaena (*30*): Notably, maternal androgens can change the cellular processes within developing foetal ovaries, altering the ratio of foetal ovarian follicles to androstenedione-producing ovarian stromal cells, thereby increasing the daughter’s ability to later raise her own androgen concentrations and, in turn, masculinise both the physiology and behaviour of granddaughters. Together, these androgen-mediated organizational and epigenetic mechanisms could ensure differential inheritance of masculinised phenotypes across generations (*30*). Although the latter possibility in meerkats requires confirmation, transgenerational female competitiveness may provide one pathway to the evolution of cooperative breeding.

## Methods

### Ethics statement

Our protocols were approved by and carried out in accordance with the Institutional Animal Care and Use Committee of Duke University (Protocol Registry Numbers A171-09-06 and A143-12-05) and the University of Pretoria’s Animal Use and Care Committee (Ethical Approval Number EC074-11).

### Study population

We studied wild meerkats in the Kuruman River Reserve, South Africa, where annual mean population size was 270 animals, in 22 established clans of 4-39 animals (*19*). Our present study period, from November 2011 to April 2015, included an extended drought, during which female reproductive success tracked rainfall (*21, 53*; Fig. S1). The population, habitat, climate and general methods have been described elsewhere (*21, 27*). The animals are microchipped, individually identifiable via unique dye marks, and habituated to close observation (< 2m) and routine weighing (*19, 27*). Meerkats reach sexual maturity at about 1 year of age and both sexes disperse at around 2 years. The matriarch, who is radio collared, holds the primary power position and has a distinct morphotype that contributes to long-term stability of her status: beyond her clan surveillance and leadership roles, she is distinguishable by weight advantage, altered body proportions, increased reproductive output and older age (*15, 54, 55*). Status and life history variables were known and monitored by non-project members.

### Focal subjects and datasets

Five of the 57 unique focal dams changed status during the study, producing 31 dominant and 31 subordinate dams that carried 89 pregnancies to term, all with live births within the clan (i.e., excluding abortions and evictions: *21*). We monitored pregnancies, detectable at 3-4 weeks of gestation, using palpation, weight gain, and shape change, to determine gestational stage in real time. We later confirmed/estimated conception by backdating 70 days from the known dates of birth (*21, 23, 27*). Given field complexities, dams contributed differentially to endocrine and behavioural datasets across EP, MP, LP and PP. Endocrine data derived from 35 individuals contributing a total of 49 serum samples (generally one blood sample per pregnancy) and 36 individuals contributing a total of 156 faecal samples, with 18 individuals contributing both types of biological samples (table 1). Adult behavioural data derived from 37 unique individuals during 48 pregnancies, and totaled 1598 focal sessions or 422 hours of observation (table 2). The focal offspring were 103 unique individuals from 32 litters, representing 14 ‘dominant control’ (DC), 13 ‘subordinate control’ (SC) and 5 ‘dominant treated’ (DT) litters (table 3). Offspring behavioural data for the first six months of life totaled 5713 focal sessions or 1376 hours of observation.

**Table 1.** Metadata for the serum and faecal samples from meerkat dams.

| Sample | Female category | Subject features |  |  | Samples by stage |  |  |  | Total |
| --- | --- | --- | --- | --- | --- | --- | --- | --- | --- |
| type |  | # | Mean | Litter | EP<sup>a</sup> | MP | LP | PP<sup>b</sup> | # |
|  |  | Ss | age (yr) | #s |  |  |  |  |  |
| Serum | Dominant, control (DC) | 18 | 4.3 | 27 | 6 | 6 | 15 | [1] | 27 |
|  | Subordinate under DC | 17 | 2.2 | 22 | 5 | 8 | 9 | [3] | 22 |
|  | Total unique used | 35 | 3.3 | 49 | 11 | 14 | 24 | [4] | 49 |
| Faecal | Dominant, control (DC)<sup>c</sup> | 20 | 4.9 | 31 | (4) | 23 | 39 | 26 | 88 |
|  | Subordinate under DC | 16 | 1.5 | 18 | (5) | 8 | 23 | 19 | 50 |
|  | Dominant, treated (DT)<sup>c</sup> | 6 | 5.4 | 7 | - | - | 17 | 1 | 18 |
|  | Total unique | 36 | 3.8 | 51 | (9) | 31 | 79 | 46 | 156 |

<sup>a</sup> Too few faecal samples (in parentheses) were obtained for analysis in early pregnancy (EP), relative to mid, late, and postpartum (MP, LP and PP, respectively). These are excluded from the total numbers reported.
<sup>b</sup> Conservatively, females were not typically captured for blood sampling during early PP, providing too few serum samples [in brackets] for analysis. These were also excluded from the total numbers reported.
<sup>c</sup> All endocrine data obtained pre-implantation from DT subjects contributed to the data from DC animals.

**Table 2.**
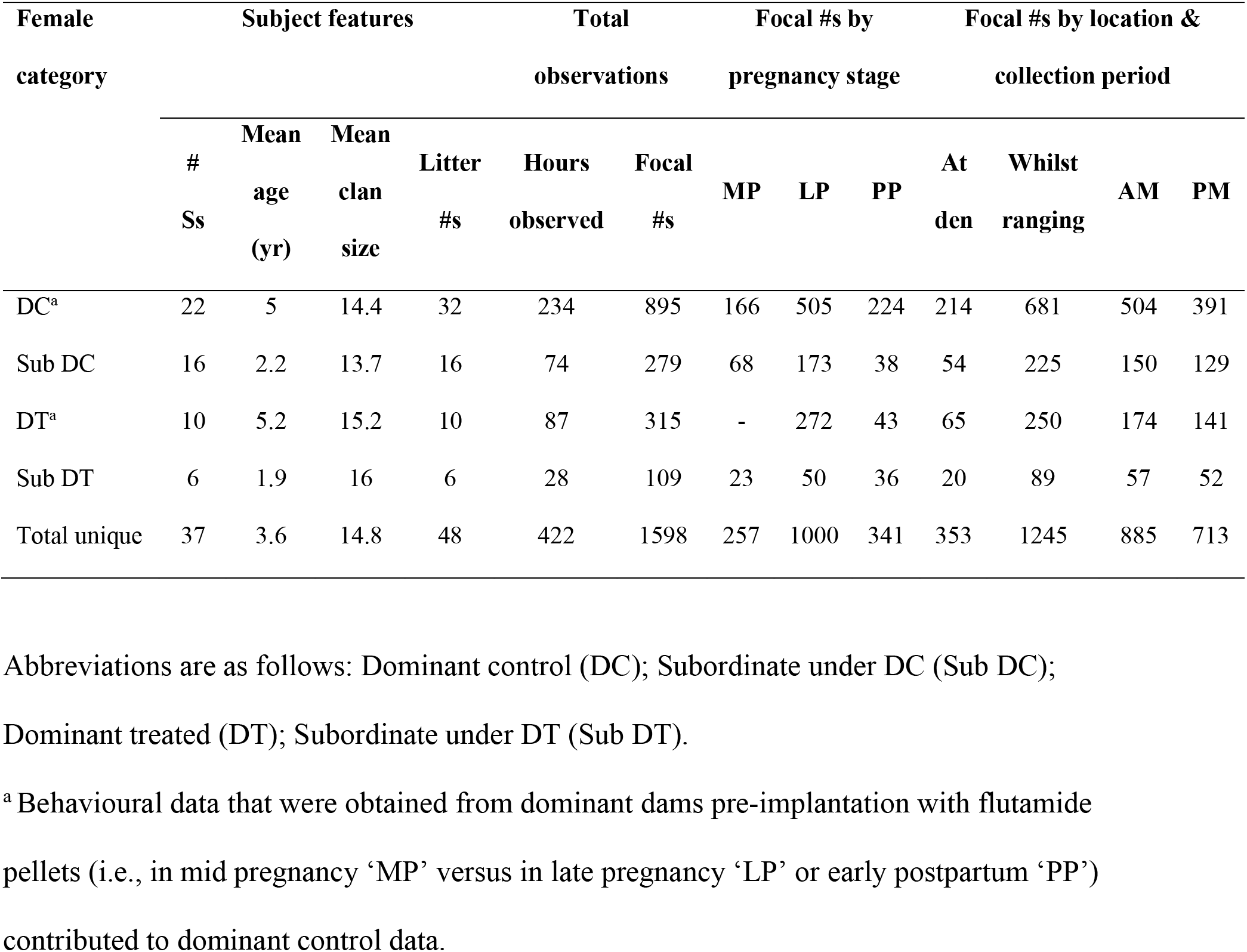
Metadata for the focal behavioural observations of meerkat dams.

**Table 3.**
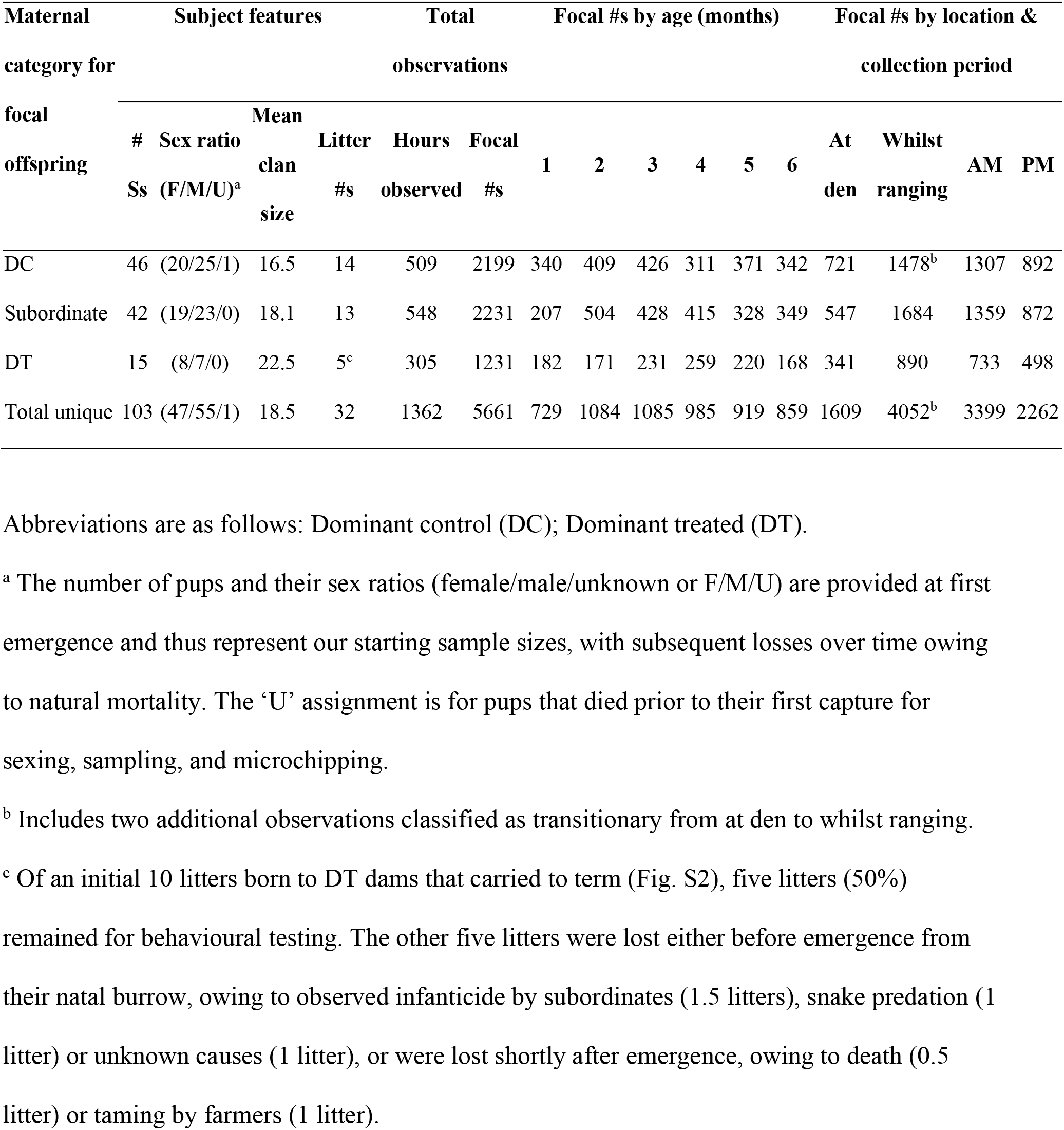
Metadata for the focal behavioural observations of meerkat offspring.

### Biological sample collection

As previously described (*19, 35*), we individually captured a target female upon emergence from the den (an underground burrow system), gently picked her up by the base of her tail, placed her into a cotton sack and anaesthetized her with isoflurane (Isofor; Safe Line Pharmaceuticals, Johannesburg, South Africa) in oxygen, using a vehiclemounted vaporiser (*56*). We used a 25G needle and syringe to first draw 0.2-2 ml of blood from the jugular vein. Thereafter, we performed additional procedures (e.g. morphological measurements) for purposes beyond the present scope. Once recovered from anaesthesia, the female was returned to her clan (20–30 min post-capture) and closely monitored throughout that and the following day, both by project and non-project personnel, including veterinarians. Behavioural data collection was suspended during this time.

We transferred the blood to a serum separator tube (Vacutainer^®^, Becton Dickinson, Franklin Lakes, NJ, USA) and allowed it to clot at ambient temperature. We later centrifuged the blood at 3700 rpm, at 24°C, for 10min and stored the decanted serum, on site, at minimally −20°C. We opportunistically collected roughly half the amount of any freshly voided faeces (to avoid disrupting faecal scent marking) into clean plastic bags, immediately placed them on ice and later stored them at minimally −20°C. After transport, on ice, to Duke University (Durham, North Carolina), we stored the samples at −80°C until analysis. We processed (lyophilised, pulverised, sifted, weighed and extracted) all faecal samples within a year of collection (*19*).

### Endocrine assays

We used competitive enzyme immunoassay kits (ALPCO diagnostics, Salem, NH, USA), previously validated for meerkats (*19, 35*) and for sample storage time (*19*), to measure androgen concentrations. The androstenedione assay has a sensitivity of 0.04ng/ml using a 25-μl dose, with intra- and inter-assay coefficients of variation (CVs) of 5.23% and 8.7%, respectively. The testosterone assay has a sensitivity of 0.02ng/ml using a 50-μl dose, with intra- and inter-assay CVs of 7.9% and 7.3%, respectively. The fAm assay has a sensitivity of 0.2–12.5ng/ml per plate, with intra- and inter-assay CVs of 7.7% and 6.2%, respectively. Details on the cross reactivities for these assays have been published (*19, 35*). Samples were run in duplicate and re-run if the CV exceeded 10%. Those samples with concentrations greater than the upper detection limit were diluted with assay buffer to maximally 1:8; the results were multiplied by the dilution factor. Samples with concentrations below the assay’s minimum detectable limit were allocated the minimum value.

### Behavioural observation

We tracked and observed clans minimally every three days; all observation protocols limit human presence to several days/wk. For candidate dams, deemed healthy and living in mid-large-sized, stable clans, we began behavioural observations after confirming pregnancy, continuing throughout gestation and PP. For offspring, we began behavioural observations when pups were ~1 month old (when they leave the natal den and reliably join the foraging clan). Following established procedures (*35*), we used focal-observation protocols (*57*), as these are less vulnerable to detection bias than are critical-incident protocols and allow capturing detailed or cryptic behaviour, including across seasons when vegetation can differentially obscure the animals. We observed focal subjects, in randomised order, during two daily periods, one beginning in early morning, after emergence, until midday (AM), the other resuming late afternoon until the animals retired underground (PM). We recorded behaviour in real time, using CyberTracker software (version 3.263, CyberTracker Conservation) uploaded to handheld palm pilots (Palm T|X, Palm, Inc.), and paused recording whenever the focal animal was out of view. We tailored our protocols to the meerkats’ major activity patterns at different times and locales: 5-min ‘den’ focals occurred during the meerkats’ brief periods spent above ground at the den, whilst mostly sedentary and prosocial (the short duration, at either end of the day, allowed rotating through focal subjects minimally once prior to the clan’s departure from or descent into the den); 30-min ‘ranging’ focals occurred in the interim, whilst meerkats primarily foraged throughout their territory.

For adults, we mostly expanded our existing ethogram (*35*) and present data on 1) aggression (including competition over an acquired food item and other high-intensity aggression or ‘HIA’; the latter comprises the former), 2) scent marking (including frenzied, body-rubbing of the environment, overmarking, defecation and urination), 3) prosociality and 4) submission, and newly added 5) ‘nearest-neighbor’ (N-N) associations (as a proxy of an ego network). For categories 1, 3 and 4, we recorded the partner and directionality of behaviour (i.e., initiator versus recipient). For category 5, every 2-3 minutes or whenever partners changed, we recorded the identities of meerkats within 10 cm or 2 m of the focal dam (for ‘den’ and ‘ranging’ focals, respectively). For offspring, we present on category 1 (with the addition of targeted beg, block approach, growl and chatter). We omitted from analysis any focal in which a disrupting ‘interclan interaction’ occurred. We assessed intra- and inter-observer reliability as before (*35*), with scores exceeding 87%; for the much longer span of offspring observations, we also confirmed that including observer identity as a random variable in the analyses had no significant impact on the findings.

### Antiandrogen treatment

*Procedural validation in males.* We planned and conducted the experimental manipulation on dominant dams only, as their reproductive success is most reliable. To minimize the number of clans affected by our protocols, as well as any potential impact of treatment on animal welfare or clan dynamics (that could potentially influence the long-term collection of normative life-history data), we had to avoid implanting matriarchs with sham pellets. Therefore, we first conducted a comparable manipulation (from Feb 2011 to Jan 2012), using the capture procedures described above, in three groups of adult, subordinate male meerkats: a flutamide-treated (~15 mg/kg/day) group, receiving 1 pellet (300 mg, Innovative Research of America, Sarasota, FL), a sham-treated group receiving 1 sham pellet of the same matrix, and untreated controls receiving no pellet (*35*). As added precaution, we brought in fulltime veterinarians, licensed in South Africa, specifically to perform the implants, as well as to monitor animals before, during and after treatment (in both studies). Other than procedures associated with pellet insertion (as described below), all capture, anaesthetic and sampling methods across subjects were identical. In males, we found (a) no negative health effects in any group, (b) no differences between sham and control groups (their health and behavioural patterns were identical), and (c) significant differences between both of the control groups relative to the flutamide-treated group, showing the predicted changes in behaviour owing to androgen receptor blockade.

### Application to females

Relying on these validated field methods (*35*) we proceeded with non-implanted control females (see below). We, thus, treated 11 dominant dams with the androgen receptor blocker, flutamide (~15 mg/kg/day), targeting the estimated last 21 days (LP) of their 70-day gestation (Fig. S2). The on-site veterinarian provided each experimental animal with a subcutaneous injection of a non-steroidal, anti-inflammatory painkiller (either 0.2– 0.3 mg/kg meloxicam, Metacam, Boehringer or 0.1 ml of metacam, plus 0.1 ml of lentrax or other long-lasting penicillin, depending on availability). Under sterile procedures, the veterinarian used a scalpel to make a 1-2 cm dorsal skin incision between the shoulder blades, blunt dissection to create a small subcutaneous pocket, forceps to insert two 21-day release flutamide pellets (as dominant females are twice the weight of adult, subordinate males), dissolvable material (Vicryl) to suture the incision and surgical glue to cover the suture for added protection.

Recovery and release of all dams was as described above (see Biological sample collection); however, treated females were monitored more extensively and for longer periods than is routine. One treated female (9%) aborted and was excluded from the study; by comparison, 21% of the contemporaneous dominant controls aborted (*21*). Given the health costs of natural androgens to dominant female meerkats (*50, 51*), flutamide-treated dams were potentially less at risk than their control counterparts. A few subjects received antibiotic ointment to treat minor infections at the suture site, but these infections typically occurred after parturition (i.e., after the cessation of focal observations) and had no noticeable effects on behaviour. Otherwise, as before (*35*), no adverse effects of treatment were detected; dominant females maintained their status and continued to fill their matriarchal role (as expected for this short treatment window).

Owing to difficulties assessing pregnancy stage in real time (*21*), particularly under drought conditions, treatment periods ultimately extended into PP, so we matched the LP+PP time span in controls (Fig. S2). Overshooting treatment was far preferable to undershooting, given that the critical period of foetal behavioural masculinisation could well occur perinatally, through a combination of androgen transfer via the placenta and maternal milk. There were no differences by age (*t*_1,35_ = 0.34, *P* = 0.738), weight (*t*_1,36_ = 0.95, *P* = 0.349) or clan size (*t*_1,39_ = 0.39, *P* = 0.699) between control and treated dominant females.

### Analysis of endocrine data

#### Main effects of status

We ran three models in R (version 3.4.3: *58*) to examine the relationship between androgens and status in control females across pregnancy. We analysed serum concentrations using GLMMs in the MASS package (version 7.3-47: *59*). We used a Gamma error distribution and log link function, accounting for females sampled across multiple pregnancies by including individual identity as a random factor. We analysed faecal concentrations using LMMs in the lme4 package (version 1.1-21: *60*). After log transformation, the response variables conformed to the normal distribution, so we used a Gaussian error distribution with an identity link function. Because we obtained multiple faecal samples per pregnancy, we included litter identity nested within individual as a random factor.

#### Main effects of trimester

To test if androgen concentrations changed with pregnancy stage, we included the interaction between status (dominant, subordinate) and stage (EP, MP, LP, PP) as fixed factors in the full models, as well as female age (continuous in years; meerkat age does not covary with status for endocrine data) and total monthly rainfall (continuous in mm^2^; rainfall highly correlates with year and territory, but was the most sensitive measure of temporal quality). Owing to drought conditions, we excluded clan identity from most analyses, because membership and location changed too often between pregnancies to be a reliable measure. Because we obtained all serum samples in the morning, we included collection period (AM or PM) only as a covariate in the fAm model (table S1).

Each full model included all probable independent terms and biologically relevant interactions; we then obtained minimal models by sequentially removing the least significant factors (*P*<0.05), starting with two-way interactions. We confirmed validity of the final models (full and minimal) using a forward stepwise procedure (*61*) and verified our assumptions by checking residuals for normality and homogeneity of variance. Collinearity between main effects was assessed via variance inflation factors (VIFs), all of which were < 2, in the R package “car” (version 2.1-6: *62*). Significance of fixed factors was determined through maximum likelihood estimation and likelihood ratio tests following a χ^2^ distribution. Significant interactions and pairs from three-level factors (e.g. EP, MP, LP or MP, LP, PP) were compared to each other using post hoc pairwise comparisons (LSD) in the lsmeans package (version 2.30-0: *63*). Here and throughout, statistical tests were two-tailed and, unless otherwise stated, we present means and standard errors.

#### Effect of antiandrogen treatment

Lastly, we analysed the effect of antiandrogen treatment on the fAm concentrations of dominant dams. Flutamide acts by blocking androgen receptors; however, in some species, it can also impact circulating hormone concentrations (*35*). Because treated females did not contribute serum samples, we addressed this question using faecal samples. We ran a new model (comparable to the prior fAm model, above) to test if flutamide influenced the fAm concentrations of treated versus control dominant dams. The full model included experimental condition (control, treatment), age, total monthly rainfall and collection period. Additionally, we included pregnancy stage (LP, PP) as a random factor to control for the timing of treatment. Comparable to subordinate male meerkats across the full three weeks of treatment (*35*), we detected no significant effect of antiandrogen treatment on fAm concentrations in dominant dams (LMM: χ^2^_1_ = 2.10, *P* = 0.147; Fig. S3).

### Analysis of adult behaviour

#### General social behaviour

To examine behavioural patterns in dams by 1) status, 2) treatment condition (for dominant females), or 3) matriarch’s treatment condition (for subordinate females), we analysed rates of aggression, scent marking, prosociality and submission using zero-inflated GLMMs in the glmmADMB package (version 0.8.3.3: *64*) in R (version 3.4.3: *58*). We focus on the MP-PP increase in behaviour, rather than the potential PP decrease, as behavioural changes routinely lag behind endocrine changes. We used either a Poisson or negative binomial error distribution, and used the distribution with the lowest Akaike’s Information Criterion (AIC) value for subsequent model selection, with focal duration as an offset. To account for repeated sampling (of individuals and pregnancies), we included litter nested within individual as a random factor, unless convergence failed, in which case we included only litter as a random factor, as this was the more sensitive measure.

#### Normative social pattern

First, we tested if behaviour (save submission, the direction of which depends on status) differed by status as the unmanipulated pregnancies progressed. We included the interaction between status and pregnancy stage (MP, LP, PP) in each full model, as well as clan size (continuous in number of members present during observation), total monthly rainfall and collection period. For behaviour that occurred irrespective of location, such as aggression (initiated or received) and scent marking, we included the type of focal (at den, whilst ranging) as a covariate. We did not do so for food competition (which occurred primarily whilst ranging) or prosociality and submission (which occurred primarily at the den). Owing to collinearity between age and status when comparing behaviour across classes, as assessed via VIFs (all > 2), we did not include age in these models (table S2).

#### Experimentally induced social patterns

Second, we tested if flutamide directly affected the behaviour of treated dominant dams, relative to control dominant dams (table S3), and third, if flutamide indirectly altered the behaviour of cohabiting subordinate dams, for which we compared the contemporaneous behaviour of subordinate dams whose matriarch was treated to that of subordinate dams whose matriarch was left untreated (table S6). In both cases (involving within-class comparisons), we included the dominant female’s treatment condition as a fixed factor in each full model, as well as the respective dominant or subordinate female’s age, clan size, total monthly rainfall and collection period. The type of focal was included as a covariate whenever relevant. Additionally, pregnancy stage for focal dominant (LP, PP) or subordinate (MP, LP, PP) dams was included as a random factor to control for potential gestational differences: Because the behaviour of subordinate dams remained consistent across pregnancy (Figs. 1D-F), we increased their sample size by extending this analysis to include subordinates at MP.

As in our endocrine analyses, we initially included all probable independent terms and biologically relevant interactions in the full models. A minimal model was obtained by sequentially removing terms based on AIC, with validity of the final models being confirmed using a forward stepwise procedure (*61*). Significance of fixed factors was determined through maximum likelihood estimation and likelihood ratio tests following a χ^2^ distribution, and significant interactions and three-level factors were compared using post hoc pairwise comparisons (LSD) in the lsmeans package (version 2.30-0: *63*).

#### Evictions

We tried several ways to analyse the number of evictions by dominant control versus treated females during LP (for *n* = 31 pregnancies). First, we excluded the pregnancies of dominant females when there were no other adult females present, then accounted for differences in clan size and number of adult females present (i.e., the number of potential eviction victims), the number of times the same female was repeatedly evicted, the timing for the onset of treatment (i.e., if victims had already been evicted when dominant females started treatment late), etc. We selected the mean percent of adult females evicted (out of the total number of adult females residing in the clan); nevertheless, accounting for these important variables for so few occurrences of evictions by treated matriarchs, reduced the power necessary to properly model this question.

#### Nearest-Neighbour (N-N) associations

To determine if antiandrogen treatment altered the matriarch’s social relationships (or centrality), we calculated dyadic rates of N-N associations (using scan sampling) for dominant dams and their adult clan members, during both treatment and control periods. We restricted this analysis to the four dominant females observed for more than an hour during both a treated and control pregnancy (totaling eight pregnancies); however, the sample size, based on Satterthwaite’s approximation method, reflects the number of adults associating with these dominant dams. The resultant dataset contained 110 dyads, including 43 adult female and 67 adult male partners (27 and 40 of which, respectively, were unique), representing 77 total hours of focal observation. Dominant females averaged 1.83 N-N scans per adult clan member per hour across periods (S.D. = 1.20, range = 0-6.35). We ran an LMM in R (version 3.6.1: *58*), using the lme4 package (version 1.1-21: *60*), with dyadic rates of N-N association as the outcome variable and partner sex, condition (treatment versus control), adult clan size, and indicator variables for dominant males and subordinate dams as main effects, and dominant female identity and partner identity as random effects. We calculated VIFs using the R package “car” (version 3.0-5: *62*); all VIFs were < 1.2, indicating adequate independence of predictor terms (table S5).

### Analysis of offspring behaviour

We used a combined model to examine the offspring’s rates of initiated aggression by maternal condition, including status (dominant versus subordinate control) and treatment (dominant control versus treated; table S6). We used zero-inflated GLMMs in the glmmTMB package (version 1.0.1: *65*) in R (version 3.6.3: *34*), a negative binomial error distribution, with focal duration as an offset. To account for repeated sampling, we included individual nested within litter nested within dam as a random factor, unless convergence failed, in which case we included only individual nested within litter, as this was a more sensitive measure. As in our analyses of adults, we included all probable independent terms (i.e., maternal condition, offspring sex, offspring age in months, clan size, rainfall, focal location and collection period) and biologically relevant interactions (i.e., between maternal condition and offspring age, and between clan size and rainfall) in the full model. We obtained a minimal model by sequentially removing terms based on AIC and confirmed final-model validity using a forward stepwise procedure (*60*). Fixed factor significance was determined through maximum likelihood estimation and likelihood ratio tests following a χ^2^ distribution, and significant interactions and three-level factors were compared using post hoc pairwise comparisons (LSD) in the lsmeans package (version 2.30-0: *62*).

## Acknowledgments

We thank the Kalahari Research Trust, Northern Cape Conservation Authority, and Kotze family for permission to work at the Kalahari Meerkat Project (KMP). We are grateful to B. Ashton, M. Böddeker, S. Cox, L. Garside, V. Goerlich-Jansson, I. Goncalves, E. Kabay, D. Pfefferle, A. Reyes, E. Terrade, M. Thavarajah, and D. Walsh for help with data and sample collection; to S. Bischoff-Mattson, D. Gaynor, L. Howell, M. Manser, L. Marris, J. Samson, M. Siekelova, N. Thavarajah, and the many KMP volunteers for logistical support; to J. Drewe, M. Lategan, J. Maud, S. Patterson, and C. Visser for veterinary care; and to I. Chandra, S. Epstein, M. Horowitz, and C. Southworth for assistance with behavioral data curation. This research was funded by the National Science Foundation (IOS-1021633 to C.M.D.). We relied on records maintained by the KMP, which has been supported by European Research Council Grant (No 294494 to T.C.-B.) and Swiss National Science Foundation Grant (31003A 13676 to M. Manser). Cambridge, Duke, and Zurich Universities supported the KMP during the span of this study.

## Author contributions

C.M.D. conceived of, designed, funded, and supervised the study, with input from T.C.-B., who also contributed access to the meerkat population and life-history records. C.S.D., L.K.G., and J.M. managed the field project and animal captures and, with D.V.B. and K.N.S.-K., collected samples and behavioral data and, with K.A.D.-S. and C.L.S., curated the data. C.S.D., L.K.G., K.A.D.-S. and K.N.S.-K. performed the extractions and endocrine assays. C.S.D., C.L.S., and J.T.F. analysed the data, prepared the figures, tables, and source datafiles, and with C.M.D. wrote the methods. C.M.D. wrote the manuscript, with edits from C.S.D., L.K.G., J.T.F., and T.C.-B.

## Additional Information

### Supplementary Information

accompanies this paper…

### Data and materials availability

All data necessary to evaluate the conclusions are present in the manuscript and/or the Supplementary Materials. Additional data related to this study may be requested from C.M.D.

