## Supplemental Materials for "A heritable androgenic mechanism of female intrasexual competition in cooperatively breeding meerkats"

**This PDF file includes:**

Figs. S1-S3

Tables S1 to S6

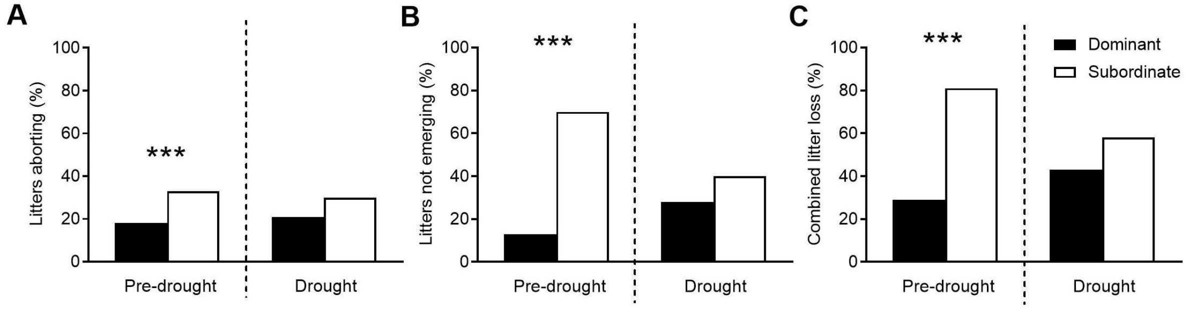


**Fig. S1. Rainfall in the Kalahari impacts status-related reproductive loss in female meerkats.** Rainfall in the Kalahari impacts insect abundance (the primary food source), with status-related consequences to reproductive loss in female meerkats. Shown for female meerkats by status are the percentages of (**A**) spontaneous abortion, (**B**) pre-emergence loss, and (**C**) both factors combined during a pre-drought period (1994-2005) and during a drought (2011-15). Relative to the earlier time frame (*31*), drought conditions were associated with both an increase in the relative, early reproductive loss of dominant females and a decrease in the relative, early reproductive loss of subordinate females, which combined to minimize female reproductive skew (*10*). *** $P < 0.001$. (These figures were created from published data; see original publications for additional statistical comparisons: *10, 37*.)

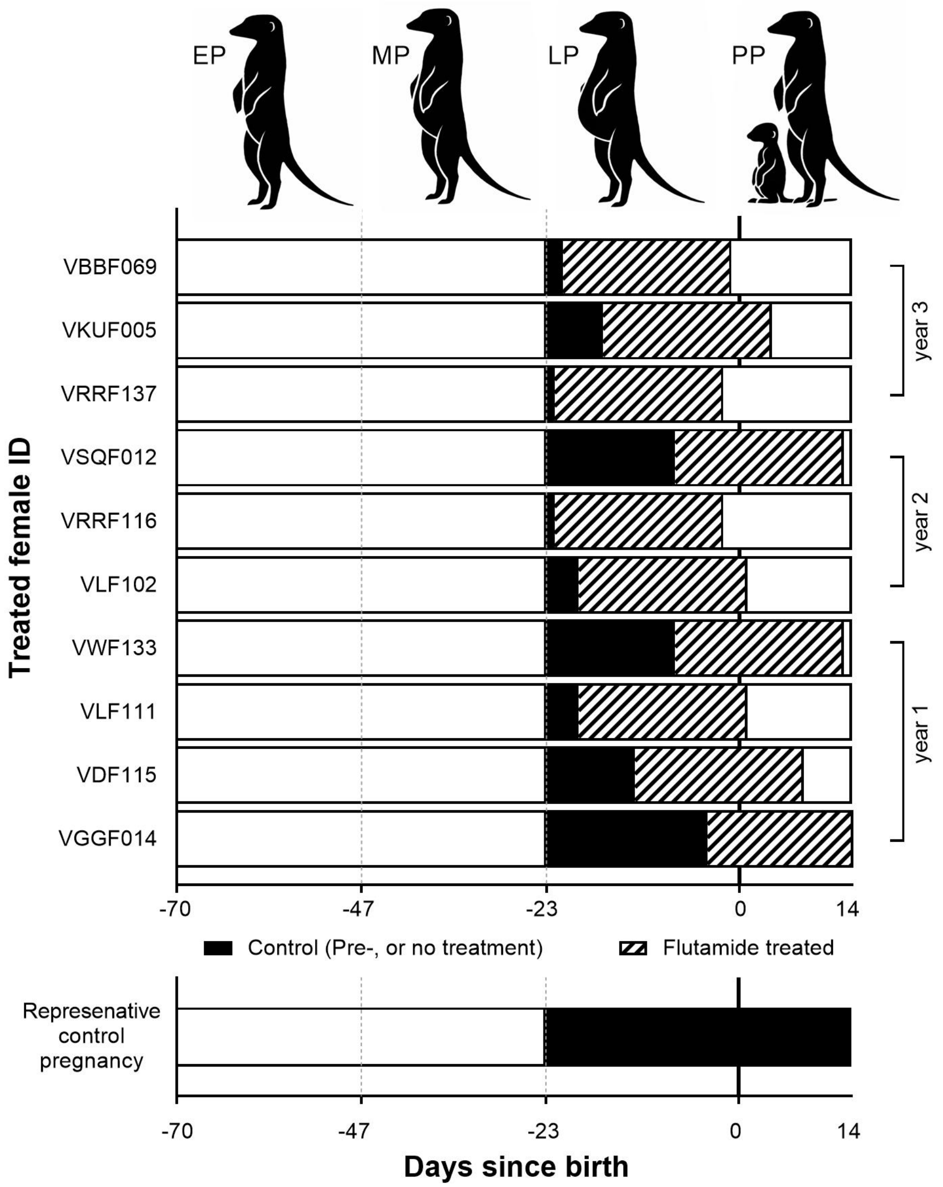


**Fig. S2. Time course of late-term antiandrogen treatment in pregnant dominant meerkats.** Across three years of study (from bottom to top), 11 dominant dams were treated with the androgen receptor blocker, flutamide; 10 of those females, shown here, carried to term. Treatment ideally targeted the last 21 days of their 70-day gestation period, although with regard to developing infants, the critical period for differentiation of brain substrates underlying behaviour is expected to be much shorter than that, on the order of days. The dark vertical line at day 0 on the x axis represents parturition, dividing pregnancy (backdated from birth) from a two-week postpartum (PP) period. Stippled lines further delineate early (EP), mid (MP) and late (LP) pregnancy. The achieved time courses of antiandrogen treatment (hatching) are shown relative to each female's own LP control period (black). EP, MP, and untreated portions of PP (white) were excluded from the analyses comparing dominant control dams to dominant treated dams. Also shown is a comparable control period extending into PP (in black) for a representative untreated, dominant dam (from a pool of 22 females and 32 pregnancies, see table 2). (Icons by S. Bornbusch.)

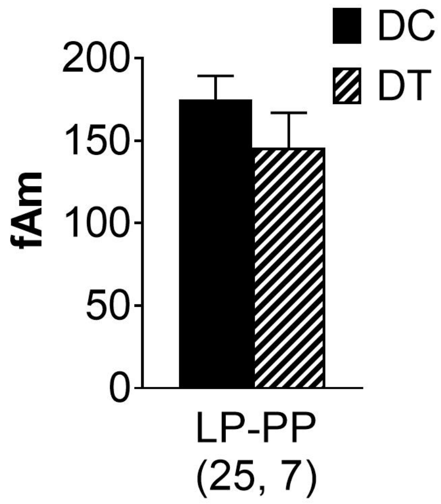


**Fig. S3. No concurrent effect of antiandrogen treatment on concentrations of faecal androgen metabolites (fAm) in meerkat matriarchs.** Values represent the mean (+ S.E.) concentrations (ng/ml) of fAm in dominant control (DC, black) versus dominant treated (DT, hatched) matriarchs across late pregnancy (LP) and early postpartum (PP). Values at the bottom of the bar graphs represent the numbers of pregnancies sampled (often multiply).

**Table S1. Status-related differences in the androgen concentrations of pregnant and postpartum meerkats.**

| Androgen | Model terms | Estimate (se) | $\chi^2$ | *P* |
|---|---|---|---|---|
| Androstenedione[a] | **Status** | **-1.765 (0.26)** | **50.49** | **<0.001** |
| | **Pregnancy stage** | | **9.50** | **0.009** |
| | ***MP*** | **-0.296 (0.32)** | | |
| | ***LP*** | **0.477 (0.29)** | | |
| Testosterone[a] | **Status** | **-1.190 (0.43)** | **6.39** | **0.011** |
| | **Pregnancy stage** | | **9.62** | **0.008** |
| | ***MP*** | **0.819 (0.37)** | | |
| | ***LP*** | **0.963 (0.33)** | | |
| Log fAm[b,*] | **Status** | **-0.412 (0.15)** | **7.60** | **0.006** |
| | **Pregnancy stage** | | **7.19** | **0.027** |
| | ***LP*** | **0.145 (0.16)** | | |
| | ***PP*** | **-0.212 (0.17)** | | |

These results derive from a minimum adequate model (MAM). All comparisons were made against the indicated levels of each factor: status = dominant; pregnancy stage = early pregnancy 'EP' (for serum values) or mid pregnancy 'MP' (for faecal androgen metabolites, fAm); collection period (for fAm) = AM. 'LP' and 'PP' indicate late pregnancy and postpartum, respectively. Bolding indicates significant terms.

[a] GLMM

[b] LMM

[*] Random effects include litter identity nested within individual (otherwise, only individual was included in the model).

**Table S2**. **Status-related differences in the behaviour of pregnant and postpartum meerkats.**

| Behaviour | Model terms | Estimate (SE) | $\chi^2$ | *P* |
|---|---|---|---|---|
| Initiate food competition[a] | **Status** | **-1.916 (0.54)** | **12.51** | **< 0.001** |
| | **Clan size** | **0.073 (0.02)** | **22.24** | **< 0.001** |
| Receive food competition[a] | **Clan size** | **0.038 (0.02)** | **4.92** | **0.026** |
| Initiate intense aggression[b,*] | **Status** | **-1.578 (0.26)** | **35.39** | **< 0.001** |
| | **Location** | **-0.453 (0.16)** | **7.67** | **0.006** |
| | **Collection period** | **-0.319 (0.12)** | **7.15** | **0.008** |
| | **Clan size** | **0.034 (0.01)** | **10.25** | **0.001** |
| Receive intense aggression[b,*] | Status | 0.1569 (0.48) | 0.03 | 0.872 |
| | **Pregnancy stage** | | **10.58** | **0.005** |
| | ***LP*** | **0.688 (0.30)** | | |
| | ***PP*** | **0.846 (0.32)** | | |
| | Location | -0.117 (0.29) | 0.06 | 0.800 |
| | **Clan size** | **0.035 (0.01)** | **5.79** | **0.016** |
| | **Total monthly rainfall** | **-0.012 (0.00)** | **8.78** | **0.003** |
| | **Status*pregnancy stage** | | **6.30** | **0.043** |
| | ***Status*LP*** | **-1.198 (0.54)** | | |
| | ***Status *PP*** | **-0.720 (0.66)** | | |
| | **Status*location** | **2.235 (0.47)** | **22.57** | **< 0.001** |
| Initiate prosociality[c,*] | **Clan size** | **0.030 (0.01)** | **3.92** | **0.048** |
| Receive prosociality[c,*] | **Status** | **-0.899 (0.23)** | **14.72** | **< 0.001** |
| | Total monthly rainfall | 0.005 (0.00) | 3.23 | 0.072 |
| Scent mark[b,*] | **Status** | **-0.474 (0.21)** | **5.31** | **0.021** |

These results derive from minimum adequate models (MAM). All comparisons were made against the indicated levels of each factor: status = dominant; pregnancy stage = mid pregnancy ‘MP’; location = den; collection period = AM. $\chi^2$ = likelihood ratio test statistic; df = 1. ‘LP’ and ‘PP’ indicate late pregnancy and postpartum, respectively. Bolding indicates significant terms.

[a] Includes focals whilst ranging only.

[b] Includes focals at den and whilst ranging.

[c] Includes focals at den only.

[*] Random effects include litter nested within individual (otherwise, only litter was included in the model).

**Table S3. Effects of mother and offspring age on offspring aggression.**

| Behaviour | Model terms | Estimate (SE) | $\chi^2$ | *P* |
|---|---|---|---|---|
| Initiate aggression[a,b,*] | **Mother** | | **13.49** | **0.001** |
| | ***Subordinate control*** | **-0.635 (0.17)** | | |
| | ***Dominant treated*** | **-1.130 (0.21)** | | |
| | **Offspring age** | **-0.364 (0.03)** | **255.98** | **<0.001** |
| | **Group size** | **0.007 (0.01)** | **8.70** | **0.003** |
| | Total monthly rainfall | -0.007 (0.00) | 0.08 | 0.772 |
| | **Location** | **2.308 (0.09)** | **639.75** | **<0.001** |
| | **Collection period** | **-0.273 (0.05)** | **27.39** | **<0.001** |
| | **Mother*offspring age** | | **24.29** | **<0.001** |
| | ***Subordinate*offspring age*** | **0.078 (0.04)** | | |
| | ***Dominant treated *age*** | **0.210 (0.04)** | | |
| | **Group size*total monthly rainfall** | **0.0004 (0.00)** | **8.24** | **0.004** |

These results derive from a minimum adequate model (MAM). All comparisons were made against the indicated levels of each factor: mother = dominant control; location = den; collection period = AM. $\chi^2$ = likelihood ratio test statistic; df = 1 for all factors except 'mother,' in which df = 2. Bolding indicates significant terms.

[a] Includes focals at den and whilst ranging.

[b] Offspring sex was not a fixed effect included in the MAM. It was not a significant predictor in the full model and was removed sequentially via AIC.

[*] Random effects include individual nested within litter nested within dam (otherwise, only individual nested within litter was included in the model).

**Table S4. Effects of antiandrogen-treatment on the behaviour of pregnant and postpartum, dominant meerkats.**

| Behaviour | Model terms | Estimate (SE) | $\chi^2$ | *P* |
|---|---|---|---|---|
| Initiate food competition[a] | **Treatment condition** | **-0.691 (025)** | **7.39** | **0.007** |
| | **Age** | **0.371 (0.06)** | **42.18** | **< 0.001** |
| | **Total monthly rainfall** | **0.008 (0.00)** | **4.28** | **0.039** |
| Receive food competition[a] | **Clan size** | **0.038 (0.02)** | **5.74** | **0.017** |
| Initiate intense aggression[b,*] | **Collection period** | **-0.304 (0.11)** | **7.51** | **0.006** |
| | Location | -0.295 (0.16) | 3.27 | 0.070 |
| | **Clan size** | **0.041 (0.01)** | **14.07** | **< 0.001** |
| Receive intense aggression[b,*] | Treatment condition | -0.306 (0.18) | 2.83 | 0.092 |
| | **Clan size** | **0.031 (0.01)** | **8.04** | **0.005** |
| | **Total monthly rainfall** | **-0.009 (0.00)** | **7.36** | **0.007** |
| Initiate prosociality[c,*] | Treatment condition | -0.450 (0.29) | 2.46 | 0.117 |
| | **Clan size** | **0.035 (0.02)** | **4.94** | **0.026** |
| Receive prosociality[c] | **Treatment condition** | **-0.507 (0.21)** | **5.73** | **0.017** |
| | **Clan size** | **0.022 (0.01)** | **5.22** | **0.022** |
| Receive submission[c,*] | **Treatment condition** | **-0.902 (0.44)** | **4.26** | **0.039** |
| | **Collection period** | **-0.629 (0.29)** | **4.81** | **0.028** |
| Scent mark[b,*] | **Treatment condition** | **-0.603 (0.23)** | **6.94** | **0.008** |
| | **Location** | **0.630 (0.25)** | **6.36** | **0.012** |
| | **Collection period** | **-0.471 (0.15)** | **10.34** | **0.001** |
| | **Clan size** | **-0.038 (0.02)** | **5.82** | **0.016** |

These results derive from minimum adequate models (MAM). All comparisons were made against the indicated levels of each factor: treatment condition = control; location = den; collection period = AM. $\chi^2$ = likelihood ratio test statistic; df = 1. Bolding indicates significant terms.

[a] Includes focals whilst ranging only.

[b] Includes focals at den and whilst ranging.

[c] Includes focals at den only.

[*] Random effects include pregnancy stage (late pregnancy and postpartum) and litter nested within individual (otherwise, only pregnancy stage and litter were included in the model).

**Table S5. Effects of antiandrogen-treatment of dominant female meerkats on her dyadic rates of nearest-neighbour associations, per adult clan member, as measured by proximity scans.**

| Behaviour | Model terms | Estimate (se) | *t*-value | *P* |
|---|---|---|---|---|
| Proximity | **Intercept** | **4.427 (0.78)** | **5.65** | **< 0.001** |
| | **Treatment condition** | **-0.602 (0.20)** | **-3.04** | **0.004** |
| | **Clan size** | **-0.158 (0.05)** | **-3.07** | **0.017** |
| | Partner is a subordinate pregnant female | -0.981 (0.51) | -1.90 | 0.060 |
| | Partner is male | -0.108 (0.25) | -0.43 | 0.670 |
| | Partner is the dominant male | 0.563 (0.43) | 1.30 | 0.199 |

These results derive from linear mixed models using matched control and antiandrogen conditions for four dominant females (i.e., $n$ = eight litters in four clans). Data were obtained during late pregnancy and early postpartum periods, from focals conducted both at the den and whilst the animals were foraging, using a scan sampling protocol.

**Table S6**. **Effects of the matriarch's treatment condition on the behaviour of pregnant and postpartum subordinate meerkats.**

| **Behaviour** | **Model terms** | **Estimate (se)** | $\chi^2$ | ***P*** |
|---|---|---|---|---|
| Initiate food competition[a,*] | **Matriarch's treatment** | **1.561 (0.61)** | **6.48** | **0.011** |
| | Collection period | 0.911 (0.59) | 2.41 | 0.120 |
| Receive food competition[a,*] | **Clan size** | **0.056 (0.02)** | **6.06** | **0.014** |
| Initiate intense aggression[b,*] | **Matriarch's treatment** | **0.940 (0.37)** | **6.55** | **0.010** |
| | Location | -1.392 (1.04) | 1.79 | 0.181 |
| | Collection period | 0.524 (0.29) | 3.31 | 0.069 |
| Receive intense aggression[b] | **Location** | **1.287 (0.40)** | **10.57** | **0.001** |
| | **Clan size** | **0.052 (0.02)** | **7.11** | **0.008** |

These results derive from minimum adequate models (MAM). All comparisons were made against the indicated levels of each factor: matriarch's treatment = control; location = den; collection period = AM. $\chi^2$ = likelihood ratio test statistic; df = 1. Bolding indicates significant terms.

[a] Includes focals whilst ranging only.

[b] Includes focals at den and whilst ranging.

[*] Random effects include pregnancy stage (mid pregnancy, late pregnancy, postpartum) and litter (otherwise, only pregnancy stage was included in the model).
